# Zone Equalisation Normalisation For Improved Alignment of Epigenetic Signal

**DOI:** 10.64898/2025.12.10.693203

**Authors:** Tom Wilson, Thomas A. Milne, Simone G. Riva, Jim R. Hughes

## Abstract

High-throughput genomic technologies have transformed our understanding of biological systems, but analysing these complex datasets remains challenging. Existing normalisation methods often struggle to accurately correct for technical variations across different experimental conditions, potentially leading to misleading biological interpretations. We introduce Zone Equalisation Normalisation (ZEN), a novel approach designed to improve signal normalisation across genomic datasets. By developing a flexible method that reduces technical noise whilst preserving biological signals, ZEN addresses critical limitations in current normalisation techniques. By applying ZEN to diverse genomic techniques, we reveal its robust performance in managing technical variations across different experimental platforms. ZEN provides researchers with a robust tool for more accurate and reliable genomic data analysis, potentially revealing biological insights that were previously obscured by technical variations.

## Introduction

High-throughput sequencing-based assays have revolutionised our understanding of gene regulation by enabling comprehensive interrogation of molecular mechanisms. Techniques such as assay for transposase-accessible chromatin using sequencing (ATAC-seq) provides insights into chromatin accessibility, which acts as a proxy to identify active enhancer and promoters (1); chromatin immunoprecipitation followed by sequencing (ChIP-seq) reveals protein-DNA interactions like transcription factors (TFs) and histone modifications (2, 3); and transient transcriptome sequencing (TT-seq) captures nascent transcription at unprecedented resolution (4–6). However, raw data generated by these technologies is inherently susceptible to technical variation that can significantly compromise the accuracy and reliability of scientific interpretation (7, 8).

Analysing these complex genomic signals therefore remains challenging. A key issue is normalisation, whereby signal tracks are scaled for direct and meaningful comparison across samples. This is a critical preprocessing step in genomic data analysis, addressing the systematic biases and technical artefacts that can potentially obscure or misrepresent genuine biological signals (9, 10). Sources of technical variation include differences in sequencing depth, experimental batch effects, sample preparation methodologies, instrument-specific technical noise, genomic content variations, and RNA degradation processes (11, 12). Many normalisation methods have been developed to address challenges with genomic data visualisation and preprocessing. For visualisation, global normalisation techniques are most commonly used, such as reads per kilobase per million mapped reads (RPKM), counts per million (CPM), bins per million mapped reads (BPM) and reads per genome coverage (RPGC), which adjust for sequencing depth by scaling raw counts to a standard reference point based on the mappable genome (13, 14). However, in practice, such methods often fail to adequately align samples across experiments as they only correct for library size (15).

For ChIP-seq, spike-in normalisation rescales signal using a stable reference derived from exogenous chromatin (16). Yet, this requires additional experimental steps and may not be comparable across experiments. For RNA sequencing (RNA-seq), housekeeping gene normalisation and quantile normalisation use internal control genes or redistribute data distributions to minimise technical variations (17, 18). More sophisticated methods like trimmed mean of M-values implemented in edgeR and DESeq2’s relative log expression normalisation use advanced statistical algorithms to account for compositional differences and systematic biases across samples (9, 19). However, these are designed for count-based data and require samples to be batch processed. Recent developments in machine learning (ML) and statistical modelling have further expanded normalisation approaches, introducing methods such as conditional quantile normalisation and probabilistic normalisation techniques that can adaptively address complex technical artefacts in RNA-seq (10, 20). Each normalisation strategy offers unique advantages and limitations, necessitating careful selection based on experimental design, data characteristics, and specific research objectives.

To address these limitations, we introduce a new normalisation method, Zone Equalisation Normalisation (ZEN), that aims to equalise variance across regions of the signal whilst being robust to extreme technical artefacts. This builds upon the normalisation method introduced by Riva *et al*. for training an ML model, REnformer, to predict chromatin accessibility (21). Furthermore, a novel approach for recreating bigWigs prior to normalisation is outlined, as well as a new way to benchmark existing methods.

## Materials and Methods

### A. ZEN workflow

Figure 1 illustrates the complete ZEN workflow, which processes genomic signals through several key stages. These include blacklist filtering, signal convolution, distribution fitting, signal zone prediction, quality filtering and signal rescaling to produce normalised bigWigs, as detailed in the following.

**Fig. 1.**
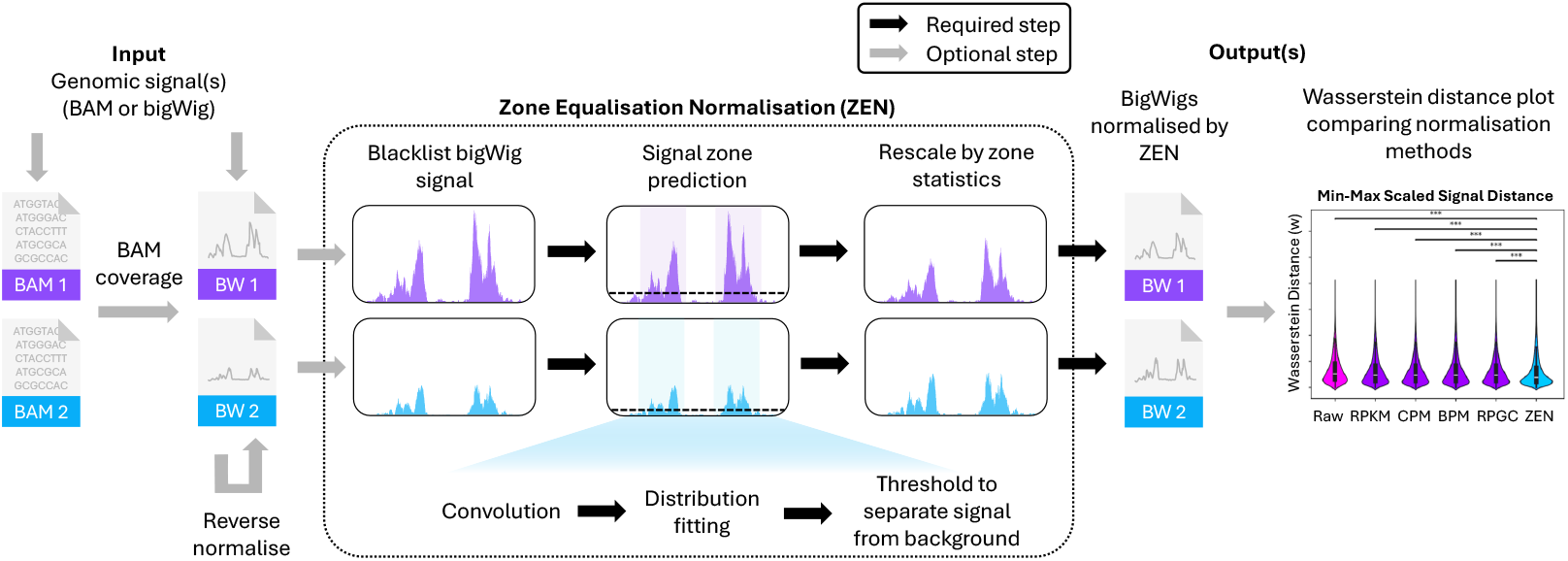
Graphical overview of ZEN. Genomic signal per sample is inputted as either BAM or bigWig format. If BAMs are given, these are converted to bigWig, while if bigWigs are directly given, these can optionally be reverse normalised. BigWig signal is then blacklisted, and signal zones predicted, which involves signal smoothing via convolution, distribution fitting and chromosome-specific thresholding to separate signal of interest from background noise. BigWigs normalised with ZEN are outputted, and alignment across samples may optionally be compared against other normalisation methods through Wasserstein distance plots.

#### A.1. Blacklisting

Considering a set of chromosomes 𝒞, the unnormalised signal is defined as 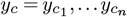, where *c* ∈ 𝒞 and *n* is the size of chromosome *c*. To begin, blacklisted coordinates are excluded from the original signal *y*_*c*_, which is read from the input BAM (mapped with deepTools bamCoverage (22)) or bigWig.

#### A.2. Signal convolution

Regions with 500 base pairs (bp) or more of consecutive zeros are masked for efficiency, creating an intermediate signal 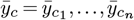. Then, 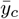 is smoothed by convolution (Eq. 1), using a normalised 301 bp triangular kernel *κ* (Eq. 2) to create the smoothed signal 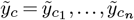. Although the kernel is customisable, *κ* was chosen as the default to fit the typical size range of regulatory element (RE) peaks. For regions shorter than 301 bp, the kernel is symmetrically trimmed to fit the region.

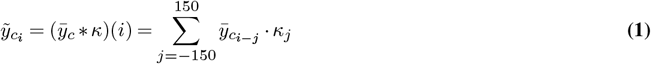

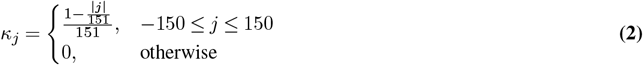

Assays such as ATAC-seq are prone to signal fluctuations, such as spikes due to random technical noise, or disruptions in alignment, like indels. Therefore, creating a smoothed signal aims to improve consistency in signal detection across genomic regions.

#### A.3. Distribution fitting

Next, zeros are removed from 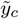 and its absolute values are log-transformed to improve symmetry and model it as a continuous statistical distribution. This creates a transformed signal, 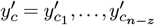, where *z* is the number of zeros for chromosome *c* (Eq. 3).

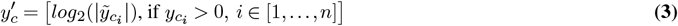

To reduce the run time and prevent over-fitting, 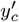 is down-sampled to *δ* positions (Eq. 4), forming 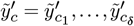, using a seed for reproducibility. By default, *δ* = 300 was chosen as this gives a good approximation of the data whilst maintaining sensitivity in Laplace distribution fitting, centred at its median and scaled by an approximation of 1 standard deviation.

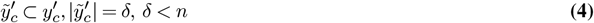

Laplace distribution is the default as it generalises well across different data types (see ‘Distribution fitting’ Supplementary section). By default, scale *V*_*c*_ (Eq. 5) is computed using the median absolute deviation (MAD) from the median: MAD(*x*) = median(|*x*_*i*_ − median(*x*)|), or mean average deviation (MeanAD) if MAD is zero: MeanAD(*x*) = mean|*x*_*i*_ − mean(*x*)|. For *d* = Laplace, constants are set as *α*_*d*_ ≈ 1.4427 and *β*_*d*_ = 1 (see Supplementary Table 1 for distribution-specific constants).

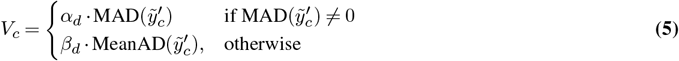

#### A.4. Signal zone prediction and quality filtering

To identify signal enriched zones, initial boundaries are estimated using the zone probability, *p* ∈ (0,1), which represents the proportion of signal to be considered background noise in the fitted distribution. For example, with the default value, *p* = 0.995 sets a cut-off between the lower 99.5% and upper 0.5% of the modelled signal. Using the fitted distribution’s percentage point function (PPF) (23), *p* is translated to a zone threshold *λ*_*c*_ (Eq. 6).

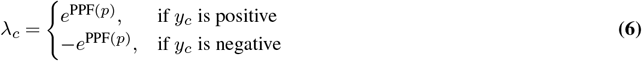

The smoothed signal, 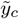, is then thresholded using *λ*_*c*_, and positions exceeding it consecutively for 35 bp or more (or below for negative signal) are set as unfiltered zones (Eq. 7). The minimum size of 35 bp aims to balance finding sufficient signal with capturing short motifs (see Supplementary section ‘Unfiltered zones minimum size’).

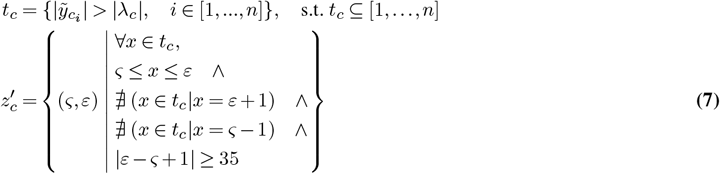

Before filtering out zones and removing any low quality signal, the estimated fragment size (Eq. 8) must first be calculated (corresponding a proxy for the value generated by a single sequenced read per chromosome), alongside the mean of non-zero signal per chromosome *µ*_*c*_ and the global non-zero mean across the genome *µ*_*g*_ are calculated (Eq. 9).

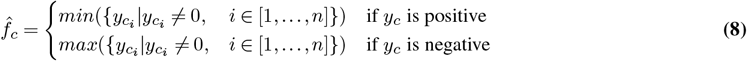

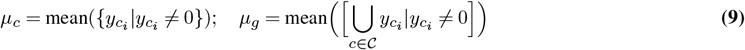

A quality threshold, *τ*_*c*_, is then defined as the maximum of the scaled estimated fragment size and the non-zero means (Eq. 10).

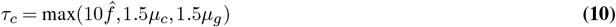

For sparse data, *τ*_*c*_ evaluates to 10 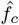. This is similar to filtering bins with less than 10 reads, as often as shown in (19, 24). For more deeply sequenced data, this may not be sufficient; consequently, *τ*_*c*_ may instead evaluate to either 1.5 times the chromosome-specific or genome-wide mean (excluding zeros). The scale of 1.5 was selected as it was found to be a good approximation of the upper limit for most values within the noise.

To refine the zones, coordinates in 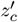 are filtered based on the original signal *y*_*c*_. Only regions where *τ*_*c*_ is exceeded for a minimum *m* bp are retained. This forms unpadded zones (Eq. 11), for which low quality signal and very sparsely mapped chromosomes are excluded. By default, *m* = 5, as this was found to be a good trade-off between filtering and retaining quality signal.

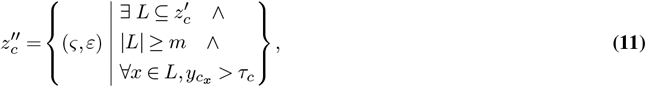

where *L* is a subset of indices between *ς* and *ε*.

Finally, padded zones are created from unpadded zones 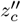, by rounding each zone to the nearest 1,000 bp and merging consecutive coordinates (Eq. 12).

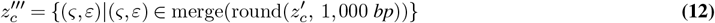

#### A.5. Rescaling signal by zone statistics

ZEN is a form of variance normalisation inspired by RNA-seq z-score normalisation (25). It adjusts the signal distribution per sample to achieve a consistent spread across sample-specific zones, thereby avoiding the need for batch processing. For each *c* ∈ 𝒞, the signal within the coordinates of each zone is extracted (Eq. 13).

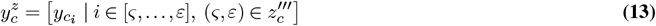

Then, extreme outliers such as technical artefacts (see Supplementary section ‘Percentile Trimming’) are excluded through removing zeros, along with the 10^*th*^ lower (*P*_10_) and upper (*P*_90_) percentiles of this non-zero signal (Eq. 14).

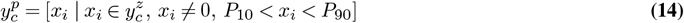

From this, the standard deviation 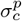 is calculated for each *c*. Then, the median 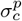 across all *c* ∈ 𝒞 are calculated as *σ*^*p*^(Eq. 15).

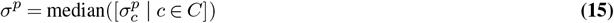

Finally, the ZEN normalised signal is created by dividing the original signal by *σ*^*p*^, therefore setting the variance as approximately 1 within non-zero signal over zones (Eq. 19). This normalised signal per sample is saved to bigWig.

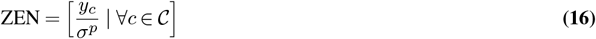

### B. Wasserstein distance distribution plots

To quantitively evaluate genomic signal alignment genome-wide across samples and replicates, we designed Wasserstein distance distribution plots which summarise pairwise signal similarity under different normalisation methods.

Signal regions were first selected and merged pairwise across samples. For punctate assays (i.e. ATAC-seq and ChIP-seq) LanceOtron peak calls were used, whereas for non-punctate assays (i.e. TT-seq) zones 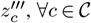 were used. To account for scale differences between normalisation methods, min-max scaling was performed between pairwise sample signals, producing two discrete one-dimensional probability distributions, *r*_1_ and *r*_2_, representing the signal distributions of these samples.

The Wasserstein distance *w* (Eq. 17) was then computed (26), where Γ(*r*_1_,*r*_2_) denotes the set of all joint probability distributions on ℝ*×* ℝ with *r*_1_ and *r*_2_ as marginals. Here, *r*_1_(*x*) and *r*_2_(*x*) specify the probability densities at genomic position *x* for each sample.

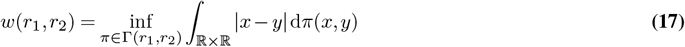

Intuitively, Wasserstein distance measures the minimal ‘work’ required to reshape one signal distribution into another. It is well-suited for comparing genomic signals as it considers both shape and spatial structure of signal distributions in a symmetric manner. Unlike correlation or point-wise comparison methods, it is robust to sequencing depth differences, technical noise and rescaling as it focuses on shape over absolute signal magnitude. This enables meaningful cross-sample and experiment comparisons recognising signals that are identical in shape and position, but differ in scale, as similar, while also capturing biologically relevant differences such as shifted binding sites, altered chromatin accessibility, or changes in transcriptional activity patterns.

### C. Data availability

For this study, we collected a diverse collection of genomic datasets to evaluate the performance of different normalisation methods across experimental contexts and cell types. For insight into chromatin accessibility and cohesin binding, ATAC-seq and ChIP-seq data were obtained from day 13 erythroid differentiation experiments. This includes bulk ATAC-seq from two donors (donors 2 and 3), each with seven replicates (reps), deposited in the Gene Expression Omnibus (GEO) under accession code GSE311157, and RAD21 ChIP-seq from three donors (donors 1, 2, and 30), each with three reps (GSE244929) (27). RAD21 donor 30 rep 2 was excluded due to low sequencing depth. Additionally, TT-seq from HeLa cells, which captures nascent RNA transcription dynamics and splicing regulation, was downloaded comparing treatment with U1-AMO to a control, each with two reps (GSE284682) (28).

Together, this collection encompasses different sequencing technologies (chromatin accessibility, protein-DNA interactions, and nascent transcription), multiple cell types (erythroid progenitors and HeLa cells), and various experimental conditions, providing a comprehensive framework to assess the robustness and applicability of ZEN across different biological contexts. All data was processed via CATCH-UP pipeline (29) with default parameters without the RPKM normalisation at the bamCoverage stage and no read extension for TT-seq. Peak calling was performed using LanceOtron, and peaks were filtered to keep those with an overall peak score *>* 0.5 (30).

### D. Code availability

All code developed for this study is open source and accessible to the research community. Source code, documentation, a tutorial and additional code to reproduce the analyses of this paper are hosted on GitHub at https://github.com/Genome-Function-Initiative-Oxford/Zone-Equalisation-Normalisation. Additionally, the ZEN method is packaged as a Python library and distributed via [PyPI](https://pypi.org/project/ZEN-norm/) and [conda](https://anaconda.org/TomMakesThings/zen-norm).

## Results

### E. ZEN improves sample alignment over functionally important loci

Normalisation is a critical preprocessing step in genomic data analysis that addresses technical variability inherent in high-throughput sequencing experiments. Differences in sequencing depth, library preparation efficiency, and instrument-specific biases can introduce systematic errors that obscure true biological signals and compromise cross-sample comparisons. Without proper normalisation, these technical artefacts can lead to false discoveries and misinterpretation of biological phenomena. Selecting an optimal normalisation strategy is therefore essential for robust and reproducible results in genomic studies, particularly when integrating data across samples, conditions, or experimental batches.

Some of the challenges of technical variability and the importance of normalisation in correcting for it are demonstrated in Figure 2, whereby raw and normalised data for three different data types are compared. For ATAC-seq, Figure 2a shows tracks from erythroid donors 2 and 6 (respectively rep 2 and rep 6) and Figure 2b shows RAD21 ChIP-seq coverage from two erythroid donors, 1 and 2 (rep 1 for both), both over the alpha-globin locus. For TT-seq, Figure 2c presents transcription from HeLa cells treated with U1-AMO (rep 1 and 2) with forward stand transcription over *CCDC127* and negative strand transcription over *SDHA*.

**Fig. 2.**
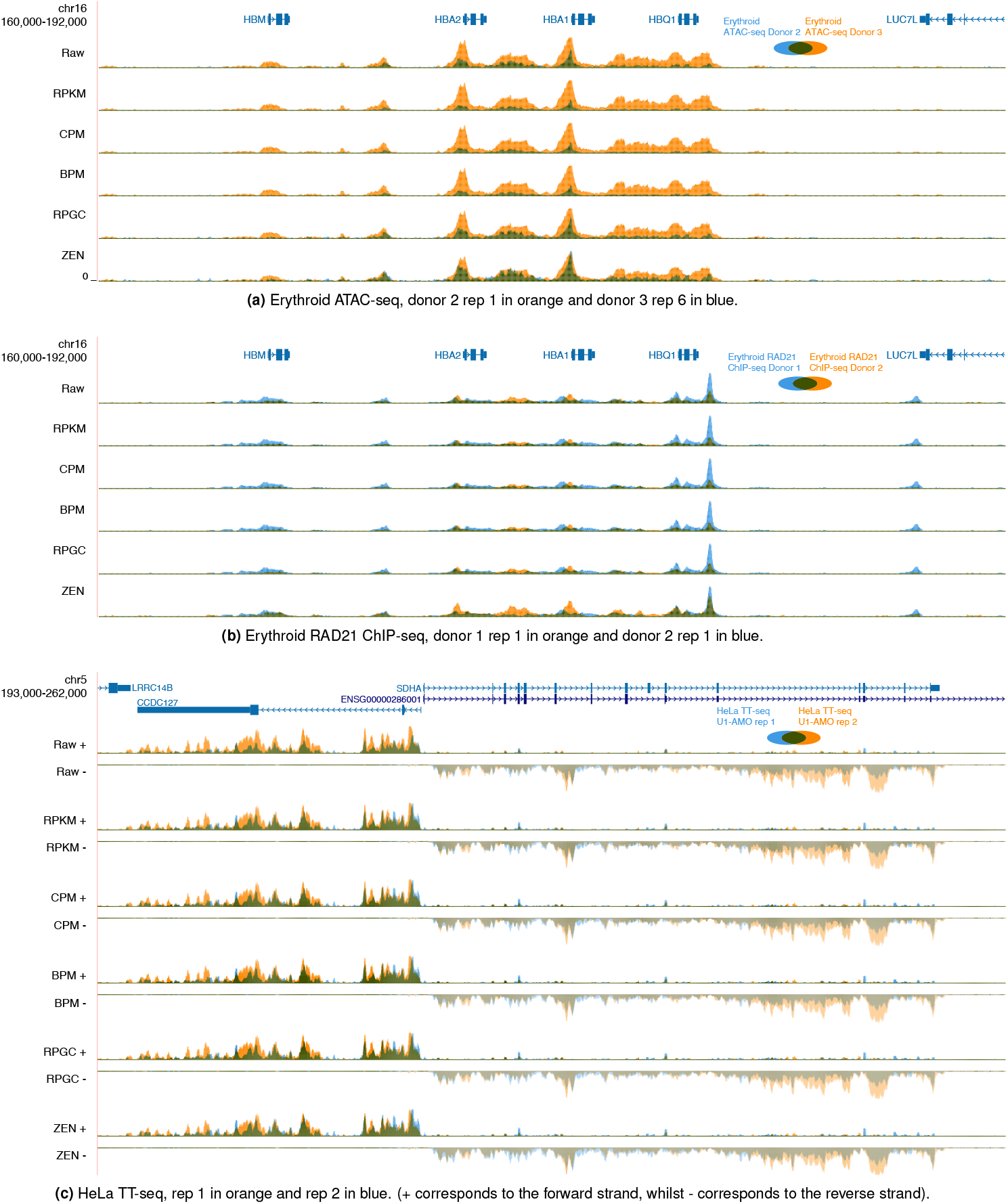
Genome Browser view of erythroid ATAC-seq and RAD21 ChIP-seq coverage over chromosome 16 (chr16:160,000-192,000) across the alpha-globin *locus* and surrounding genes (*HBM, HBA1, HBA2, HBQ1*, and *LUC7L*), respectively in (a) and (b). While (c) shows HeLA U1-AMO TT-seq over chromosome 5 (chr5:193,000-262,000) with forward strand transcription over *CCDC*127 and reverse strand transcription over *SDHA*. For each, six overlaid tracks display the same sequencing data after different normalisation methods: Raw (unnormalised read counts), RPKM, CPM, BPM, RPGC, and *ZEN*. In (a), ATAC-seq coverage from erythroid donor 2 is shown in orange and donor 3 in blue. For (b), RAD21 ChIP-seq donor 1 coverage is orange and donor 2 in blue. In (c), transcription from HeLa rep 1 is in orange, whereas rep 2 is blue. For all three, overlap between replicate signal is in green.

To benchmark ZEN, we compared it against established genomic signal normalisation techniques - RPKM, CPM, BPM, and RPGC - which adjust sequencing depth, total mapped reads, raw counts, and genome-wide coverage, respectively. Although widely used for cross-sample comparison, these conventional methods were not originally designed for that purpose (15).

We used ZEN, fitting Laplace distribution for ATAC-seq, Logistic distribution for RAD21 ChIP-seq (adjusting the zone probability to *p* = 0.999, for improved performance over the default parameters), and a left-skewed Gumbel distribution for TT-seq, as these models best capture the signal characteristics of each assay type (see ‘Distribution fitting’ Supplementary section). Selecting appropriate probability distributions (and other parameters) is critical for ZEN’s performance, as different genomic assays, cell types and states exhibit distinct properties and noise distributions. The Laplace distribution is effective at modelling the sharp, peaked signals typical of chromatin accessibility (ATAC-seq), while the Logistic distribution suits protein-DNA binding events (ChIP-seq). The left-skewed Gumbel distribution best represented the diffuse transcriptional signals in the HeLa TT-seq, but is not universally optimal for this assay. Comprehensive analyses (see ‘Distribution fitting’, Supplementary section) justify our distribution choices and validate that the selected models accurately capture the empirical data characteristics across experimental conditions and sample types ensuring optimal normalisation per experiment.

By rigorously comparing the four conventional normalisation methods with ZEN, we aim to demonstrate the method’s superior performance in mitigating technical variations, reducing batch effects and preserving biological signal integrity across diverse genomic datasets. Our analysis encompassed multiple high-throughput sequencing assays, including ATAC-seq, ChIP-seq and TT-seq to provide a comprehensive evaluation of normalisation strategies across different genomic contexts.

### F. ZEN outperforms conventional methods for genome-wide sample alignment

While ZEN improved sample align ment over key loci compared to state-of-the-art approaches, a comprehensive assessment requires genome-wide evaluation. To address this, for each dataset, we created Wasserstein distance distribution plots and across replicates using LanceOtron peaks for ATAC-seq and ChIP-seq, and zones for TT-seq. The violin plots in Figure 3 show Wasserstein distance from genomic tracks of raw (unnormalised) data, conventional normalisation methods (RPKM, CPM, BPM, RPGC) and ZEN per dataset. Each violin represents the probability density of distance values, where wider sections indicate more frequent values and Figure 3a presents distances of all pairwise comparisons between erythroid ATAC-seq samples from donors 2 and 3 (seven reps each) across LanceOtron peak calls with an overall peak score threshold *>* 0.5. Figure 3b shows the corresponding analysis for RAD21 ChIP-seq data from donors 1, 2, and 30, with all pairwise comparisons across two reps per donor. Figures 3c and 3d depict pairwise distances across all transcription zones (with padding) in TT-seq from HeLa cells, comparing U1-AMO treated and control samples with two reps each.

**Fig. 3.**
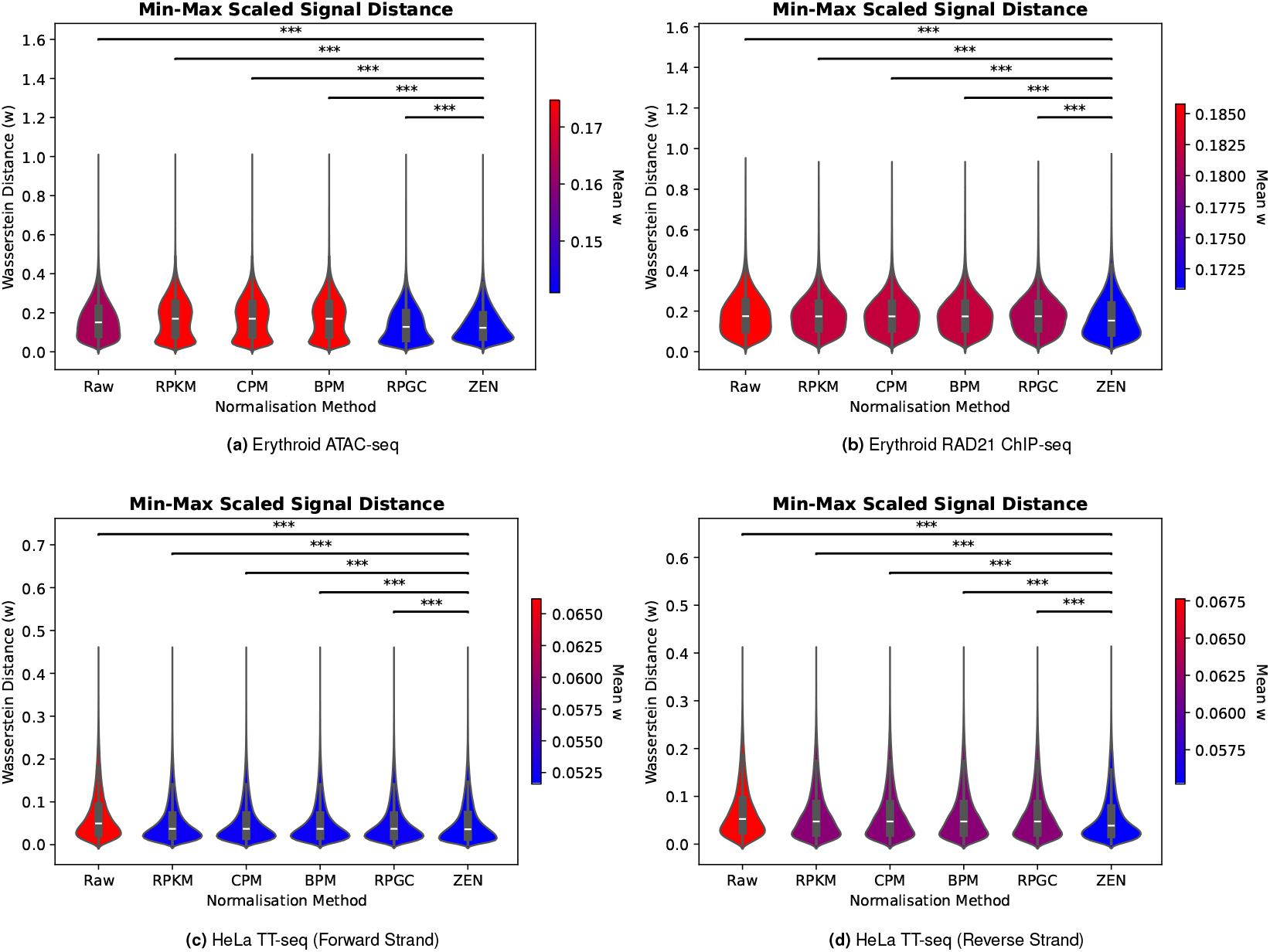
Comparing normalisation methods by min-max scaled Wasserstein distance between sample pairs across signal regions genome-wide. For (a) erythroid ATAC-seq and (b) erythroid RAD21 ChIP-seq donors, signal regions are over peaks; whereas for (c) HeLa TT-seq, signal regions are transcription zones (padded zones). Violin plots are coloured by mean Wasserstein distance, where a lower distance implies better alignment. Above each plot are p-values from t-tests comparing pairwise normalisation method distributions, where *** is p-value *<* 0.001. For ATAC-seq, the Laplace distribution was fitted with zone probability *p* = 0.995. For RAD21 ChIP-seq, Logistic was fitted with zone probability *p* = 0.999. For TT-seq, left-skewed Gumbel was fitted with zone probability *p* = 0.995.

Across all four panels in Figure 3, ZEN consistently achieves the lowest mean distances over all datasets (ATAC-seq, RAD21 ChIP-seq, and TT-seq). In every comparison between ZEN and both the unnormalised and conventional methods, distances are statistically significant (p-value *<* 0.001), as determined by pairwise t-tests, demonstrating its superior performance at aligning genomic signals between samples. Its more compact distributions, compared to the wider and more variable distributions of other normalisation methods, validate that ZEN provides substantially more stable and precise genomic track comparisons. This translates to improved reproducibility in cross-sample analyses, as evidenced by the reduced variance in distance measurements, and consistent alignment quality. This is particularly crucial for large-scale genomic studies, where consistent normalisation performance across diverse sample conditions is essential for reliable biological interpretation.

Another interesting observation is that conventional methods show relatively similar mean distances to each other within each dataset. This is likely as they all globally rescale tracks indiscriminately across signal and background, rather than basing normalisation on biologically relevant signal regions. This may also explain why they do not perform as well as ZEN, sometimes achieving only marginal improvements over unnormalised data. However, one exception was observed for the erythroid ATAC-seq, whereby RPGC performed better than the unnormalised data, while RPKM, CPM, and BPM performed worse.

Additionally, mean distances varied between assay types, with ATAC-seq and RAD21 ChIP-seq displaying means around 0.1 − 0.2, whilst TT-seq data show much lower values (around 0.02 − 0.05). This may reflect different sequencing depths, different levels of inter-sample variability between assay types or be a consequence of wider regions being compared for TT-seq. As ZEN consistently outperformed across these diverse genomic contexts (chromatin accessibility, protein binding, and transcription), this demonstrates its broad applicability and robust performance across different biological systems. Such ability to reduce technical signal variability, whilst preserving the underlying biological signal relationships, are crucial for downstream analyses.

## Conclusions

Our comprehensive analysis underscores the critical role of robust normalisation strategies in high-throughput genomic research. ZEN provides a powerful and adaptive approach that outperforms conventional normalisation methods, such as RPKM, CPM, BPM, and RPGC, across diverse genomic datasets, including ATAC-seq, ChIP-seq, and TT-seq. By equalising variance across genomic regions, whilst minimising the influence of technical artefacts, ZEN consistently improves signal-to-noise ratio, enhances cross-sample comparability, and enables more reliable detection of biologically meaningful signals.

Beyond methodological refinement, choice of normalisation method can fundamentally alter scientific interpretation. Inappropriate normalisation can lead to spurious conclusions, obscure biological signals, or introduce artefacts that compromise research integrity. ZEN addresses these challenges, offering researchers a robust and generalisable framework for genomic data preprocessing - an increasingly critical need as sequencing technologies advance and genomic assays grow in complexity.

To extend its reach, we developed a reverse normalisation tool, detailed in the ‘Reverse normalisation’ Supplementary section, allowing users to work with bigWig files that have already been processed using conventional methods. This tool reverses existing linear normalisation transformations, enabling ZEN to be run on pre-processed datasets, therefore greatly expanding its utility. Whether working with newly generated data or public repositories, ZEN provides a comprehensive solution allowing analyses that would otherwise be confounded by incompatible normalisation strategies.

In summary, this work shows that in genomic research, high-quality data preprocessing can be as crucial as experimental design. By providing a sophisticated and tailored normalisation approach, ZEN empowers researchers to uncover deeper and more reliable insights into the molecular mechanisms underlying biological systems. This versatility ensures that improved signal alignment and reduced technical variability can benefit the entire spectrum of genomic research, from individual laboratory studies to large-scale meta-analyses, ultimately advancing our collective understanding of genomic function and regulation.

## ACKNOWLEDGEMENTS

T.W is supported by the MRC grant (MC_ST_00029). T.A.M is funded by the MRC grants (MC_UU_00016/6 and MC_UU_00029/6). S.G.R. is supported by the MRC grant (MC_UU_00029/3). J.R.H. is supported by the Wellcome Trust grants (225220/Z/22/Z and 106130/Z/14/Z) and the MRC grant (MC_UU_00029/3).

## Declaration

J.R.H. is a co-founder and director of Nucleome Therapeutics and provides consultancy to the company. T.A.M. is a shareholder in and consultant for Dark Blue Therapeutics. These authors declare no other financial or non-financial interests. The remaining authors declare no competing interests.

## Supplementary Note 1: Supplementary

### A. Additional data

To ensure ZEN is designed to generalise across assay types, we tuned parameters using additional ATAC-seq, ChIP-seq and TT-seq datasets. These include ATAC-seq of brain differentiation from the H1 cell line to forebrain, midbrain and hindbrain, with three replicates per stage (GSE286146), and single cell ATAC-seq (scATAC-seq) bigWigs of 222 cell types from the human tissues [CATlas portal](https://decoder-genetics.wustl.edu/catlasv1/catlas_downloads/humantissues/Bigwig/) (31). Erythroid CTCF ChIP-seq from Donors 1, 2, and 30 is from the same study as the RAD21 ChIP-seq (GSE244929) (27), while RNA Polymerase II Ser2P (Pol II) ChIP-seq from the HEL leukaemia cell line was included with three replicates (GSE84157). TT-seq was obtained of the A-375 melanoma cell line comparing a CDK13 mutant clone R860Q with a control (GSE223888) (32), as well as HEK293T embryonic kidney cell line with four replicates (GSE218127) (33). In addition, we also downloaded H3K27ac ChIP-seq from ENCODE of E14 mouse embryonic stem cell (E0 mESC, GSE31039) and embryoid body (EB, GSE120376) (34) to demonstrate a problematic technical artefact.

All raw data was realigned using the CATCH-UP pipeline with default parameters without RPKM normalisation. Whilst for the pre-normalised CATlas bigWigs, the MACS2 Signal Per Million Reads (SPMR) (35) normalisation was reversed to create bigWigs of raw signal using the method described in the Supplementary ‘Reverse normalisation’ section.

### B. ZEN normalisation workflow

#### B.1. Distribution fitting

To determine the best distribution for ATAC-seq, ChIP-seq and TT-seq, distributions with two parameters were fitted per *c* ∈ 𝒞 for each sample in each dataset. These parameters are the location *L*_*c*_ (the shift of the distribution) and scale *V*_*c*_ (the statistical dispersion). The set 𝒟 of distributions includes: Normal, Logistic, Laplace, Gumbel (left or right) and Cauchy. For each distribution *d* ∈ 𝒟, three types of (*L*_*c*_,*V*_*c*_) parameters (referred to as parameter type) were fitted. The first infers (*L*_*c*_,*V*_*c*_) by maximum likelihood for *d* using SciPy (36). The second sets 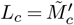 and the third sets 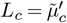, where 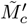 is the median and 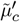 the mean of 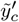 retrospectively. For these, scale *V*_*c*_ is determined by Eq. 18 using constants specified in Table 1.

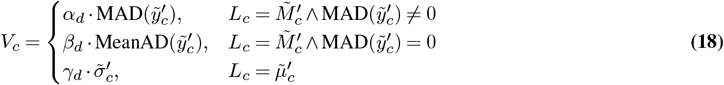

Inspecting Quantile-Quantile (Q-Q) plots (Figure 4) and histograms (Figure 5) of transformed signal 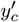, distributions for all ATAC-seq and ChIP-seq were relatively symmetrical and showed much lighter tails than the Cauchy distribution. Therefore, Cauchy and Gumbel were not deemed appropriate for these datasets. In contrast, some TT-seq datasets, such as the HeLa and HEK23T TT-seq, displayed skewed distributions, whereas the A-375 TT-seq was more symmetrical.

**Fig. 4.**
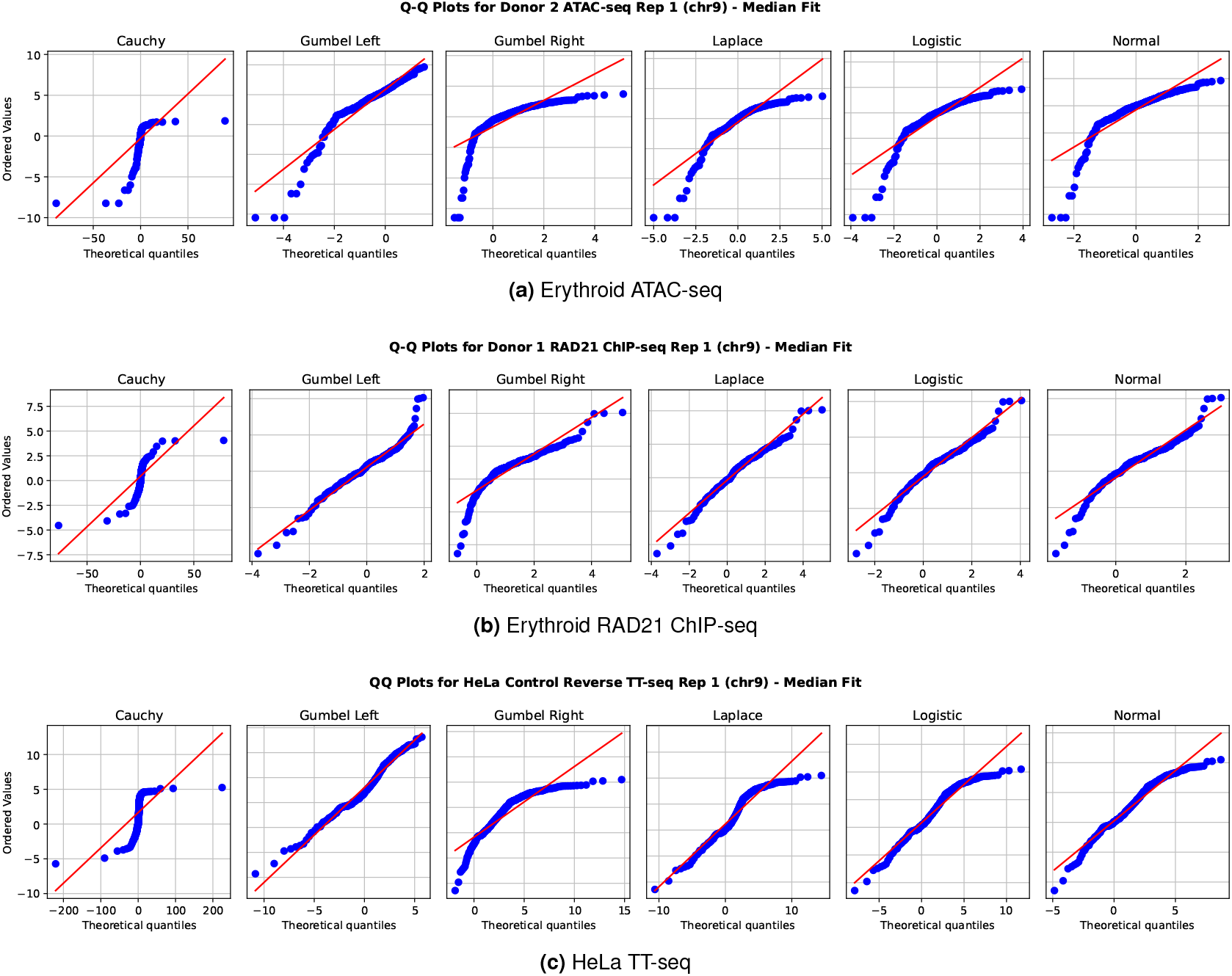
Q-Q plot per distribution fitted with median parameters to transformed signal 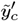 for (a) erythroid Donor 2 ATAC-seq rep 1, (b) RAD21 ChIP-seq rep 1 and (c) Hela TT-seq rep 1. Red lines are theoretical distributions, while blue dots are transformed signal.

**Table 1.** Constants for deriving the scale of a distribution *d. α*_*d*_ and *β*_*d*_ are used if *d* is centred using the median and *γ*_*d*_ if centred using the mean. Each *d* sets dispersion differently, so *α*_*d*_ is set using the distribution’s definition of MAD, *β*_*d*_ from its MeanAD and *γ*_*d*_ from its SD with several exceptions: Cauchy distribution variance is undefined, so *α*_*d*_, *β*_*d*_ and *γ*_*d*_ are set as one; Logistic distribution MAD is undefined, so *alpha*_*d*_ is one; Gumbel distribution *α*_*d*_ was derived from numerical approximation by A. Akinshin (37–39).

| $d$ | Scale Constants | | |
| --- | --- | --- | --- |
| | $\alpha_d$ | $\beta_d$ | $\gamma_d$ |
| Normal | 1.4826 | 1.2533 | 1 |
| Laplace | 1.4427 | 1 | 0.7071 |
| Logistic | 1 | 0.7213 | 0.5513 |
| Cauchy | 1 | 1 | 1 |
| Gumbel | 1.3036 | 1.6654 | 0.7797 |

**Fig. 5.**
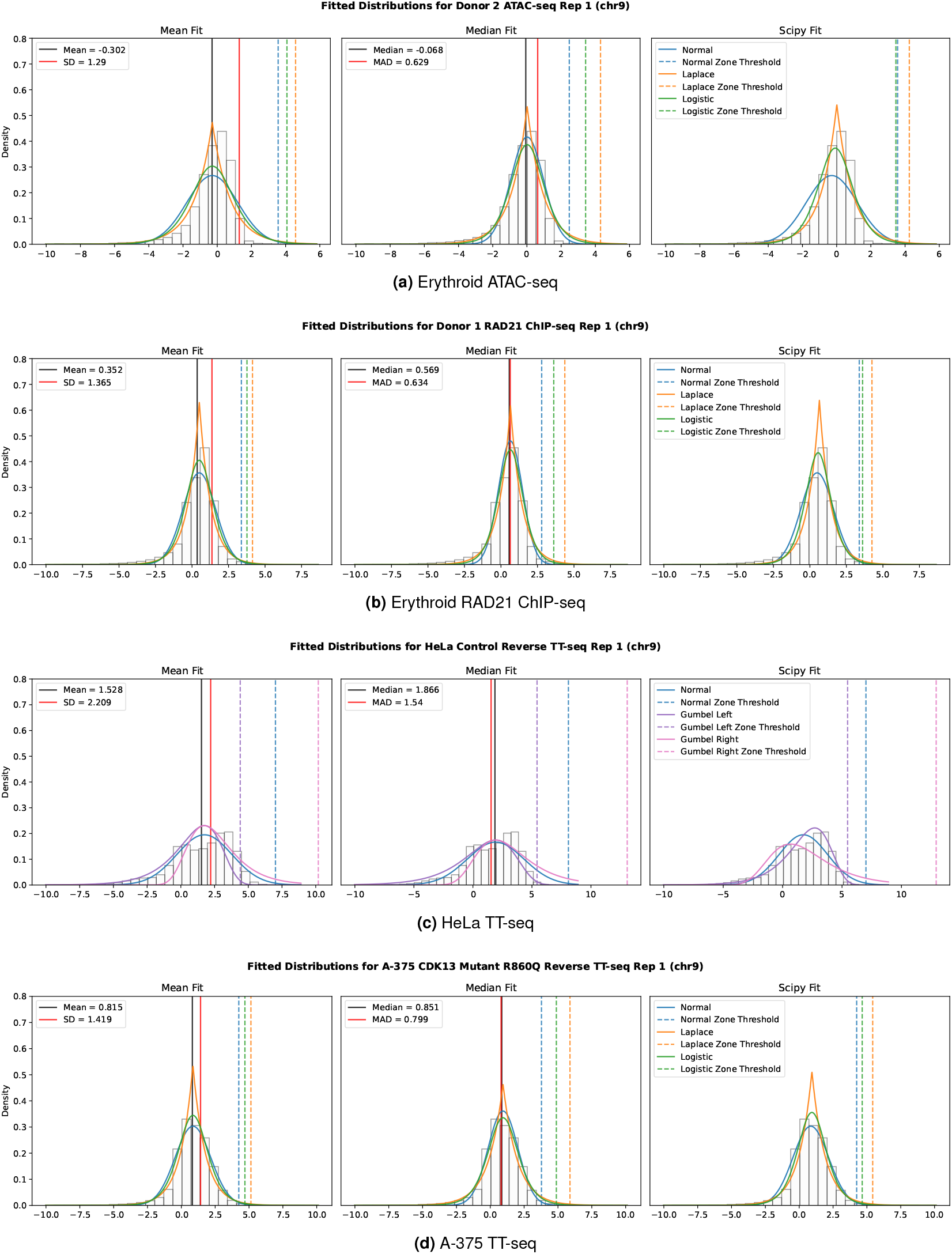
Histograms of transformed signal 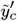 and distributions fitted with the mean (left), median (centre) and SciPy (right) parameters. For mean and median fits, black lines show a distribution’s location and red lines are the value the scale is derived from prior to adjustment. Dashed lines represent the CDF’s position for 0.995 zone probability.

The general performance of each distribution and parameter type per assay was measured by calculating the mean Kolmogorov Smirnov (KS) statistic *D*_*δ*_ for each chromosome per sample across multiple dataset and visualising results as box plots (Figure 6). Across the ATAC-seq data, the best distribution was Laplace with median parameters, while for ChIP-seq and TT-seq, Logistic with SciPy parameters performed best. Therefore, we selected Laplace for our erythroid ATAC-seq analysis and Logistic for the erythroid RAD21 ChIP-seq. However, we decided to select Gumbel with SciPy parameters for the HeLa TT-seq due to the specific properties of this data.

**Fig. 6.**
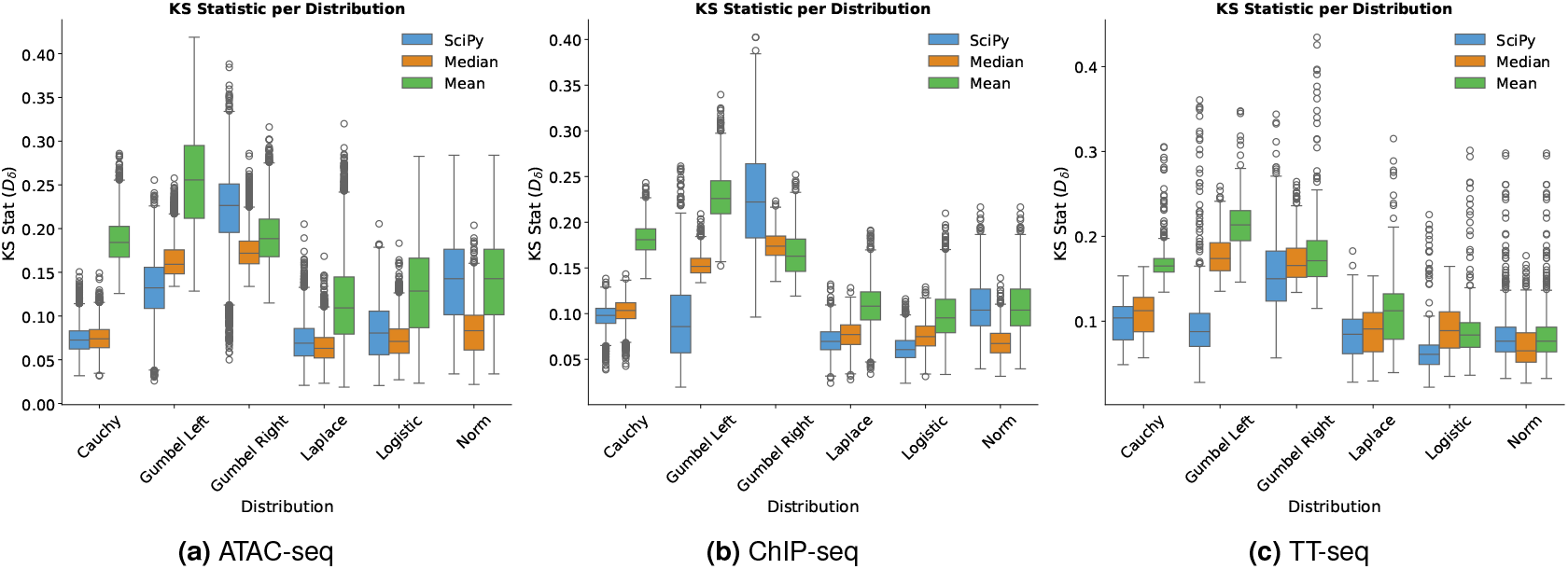
KS test results of each distribution and parameter type fitted prior to signal zone prediction. For (a), tested datasets include erythroid Donor 2 and 3 ATAC-seq, H1 brain ATAC-seq and CATlas scATAC-seq. For (b), ChIP-seq datasets include Erythroid RAD21, Erythroid CTCF and HEL Pol II. For (c), TT-seq dataset include HeLa, A-375 CDK13 (mutant and control) and HEK293T.

The zone probability parameter (*p*) may need adjusting depending on the distribution’s tail behaviour. For example, the default *p* = 0.995 for the heavier-tailed Laplace distribution produces a higher zone threshold than the Logistic distribution, as heavier tailed distributions have more probability in the extremes (see Figure 5). To maintain a comparable threshold, we therefore increased the zone probability to *p* = 0.999 when using Logistic for the RAD21 ChIP-seq.

#### B.2. Unfiltered zones minimum size

Most TF motifs — short DNA sequences that serve as binding sites for regulatory proteins - are between 5-20 bp (40). However, short read sequencing technologies keep fragments at least 50 bp in length (41), while the typical fragment size for ATAC-seq and ChIP-seq is between ∼100-600 bp (42, 43). This means that enriched peaks often span at least 100 bp. Therefore, a minimum of 35 bp consecutively exceeding the zone threshold *λ*_*c*_ was chosen to capture regions that are longer than individual motifs yet shorter than full peak widths. This ensures that the detected regions likely represent meaningful biological signals, including motifs and their immediate context, while allowing for minor local signal fluctuations within broader peaks.

For example, Figure 7 presents a genome browser view of unnormalised ATAC-seq from erythroid Donor 3 (rep 6) over the functionally important *NPRL3 locus*. This demonstrates how open chromatin peaks correspond with regions enriched in short motifs, such as *KLF4, NFX1, SP4* and *CTCF*. The zone threshold line is the cut-off by which the unnormalised signal must consecutively exceed for 35 bp or more to be classed as signal over noise. This shows how peaks are successfully separated from the background noise.

**Fig. 7.**
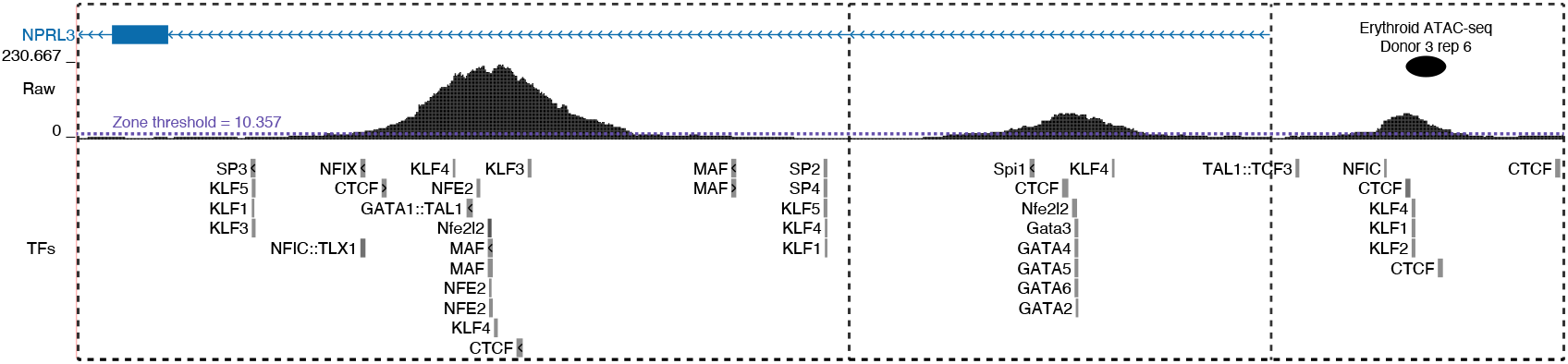
Chromatin accessibility signal tracks from erythroid Donor 3 (rep 6) without normalisation, displaying three open chromatin peaks over globin enhancers. In order, these are enhancer R2 (chr16:112,529-114,645), enhancer R3 (chr16:119,471-120,625) and enhancer R4 (chr16:142,840-143,636). The zone threshold line at ∼ 10.357 is the cut-off which signal must consecutively exceed to be considered signal rather than noise. Lower annotations show selected TF binding sites (JASPAR), representing short DNA sequences recognised by specific TFs.

#### B.3. Percentile Trimming

Even after removing blacklisted regions, the original signal *y*, some technical artefacts may remain. For example, a region with abnormally high signal can be seen in the HeLa TT-seq within chromosome 21 (Figure 8). This can occur across assay types and species. For example, Figure 9 shows an artefact in mouse embryoid body (EB) H3K27ac ChIP-seq. When calculating statistics from signals within zones, trimming 10% from either tail of the signal therefore aims to make the calculations more robust by reducing bias from these regions.

**Fig. 8.**
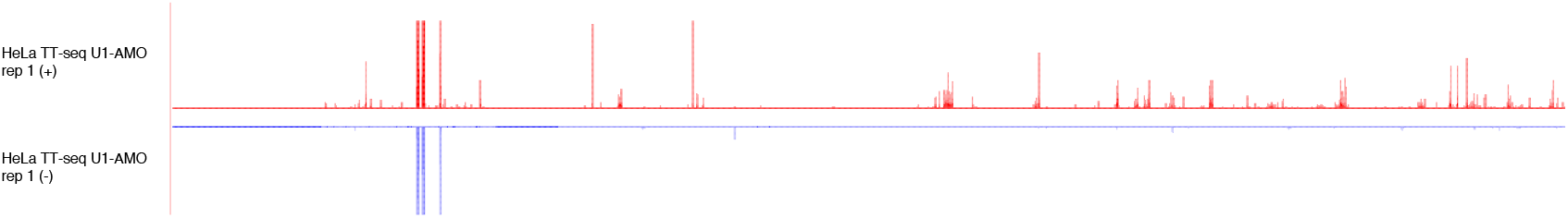
Genome browser view of HeLa TT-seq data from U1-AMO treated samples, across the full length of chromosome 21. The top track (red) from U1-AMO treated samples displays typical transcriptional regions indicating transcriptionally active genes and enhancers on the forward strand. The bottom track (blue) shows that one region has extremely high valued signal, likely due to a technical artefact.

**Fig. 9.**
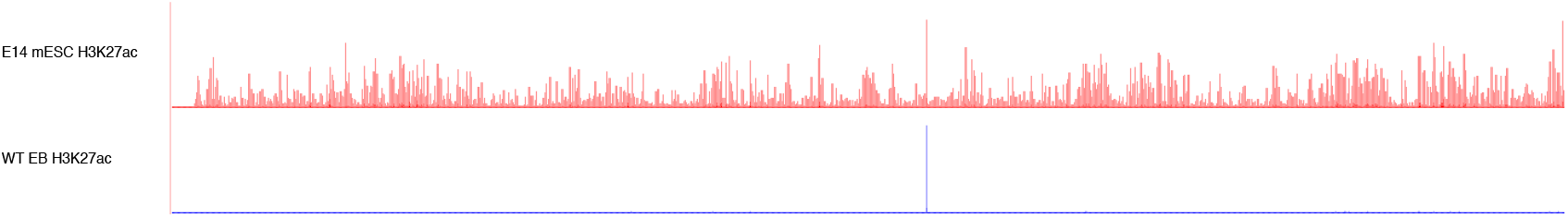
Genome browser view of H3K27ac histone modification signal across the full length of chromosome 2. The top track (red) from E14 mESC displays typical H3K27ac marking active enhancers and promoters, whilst the bottom track from WT EB shows an extreme technical artefact that has a far greater value than the rest of the signal.

#### C. Reverse normalisation

For an unnormalised signal *y* = *y*_1_,…*y*_*N*_, where *N* is the genome size, a linearly normalised signal is derived by multiplying with a scaling factor *s* ∈ ℝ^+^ (Eq. 19). Examples of such linear normalisation methods include RPKM, CPM, BPM, RPGC, and ZEN.

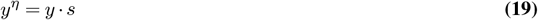

Linear normalisation can therefore be reversed using the reciprocal of *s*. But in practice, it can be difficult to recover *y* if *s* is unknown. For example, with RPKM normalisation, *s* relates to the library size (the total reads for the sample), and without access to the raw data, such as BAMs, *s* cannot be determined. However, given the high abundance of background noise in sequencing data, it is often possible to reconstruct *y* with high accuracy by estimating *s*.

Scaling factor estimation relies on the principle that the fragment size *f*, representing the contribution of a single read in *y*, is +1 if the signal is all positive and −1 otherwise. After linear normalisation, the per-read contribution of *±*1 is scaled. Therefore, the normalised fragment size *f*^*η*^ becomes *±s*, and *s* is the absolute value of *f*^*η*^(Eq. 20).

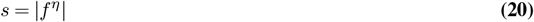

For most genomic assays, the abundance of background noise makes it highly likely that there exists at least one *i* such that 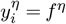. The normalised fragment size 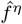 can therefore be estimated as the smallest non-zero value genome-wide (Eq. 21).

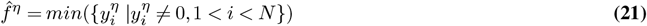

Consequently, the estimated scaling factor *ŝ* is defined as the absolute value of this estimation (Eq. 22).

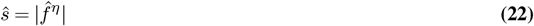

If *ŝ*= 1, this implies either *y*^*η*^ = *y* or the normalisation was non-linear. To distinguish between these cases, the signal is checked to test whether 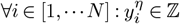. If this holds, then the signal has likely not been normalised. Otherwise, a normalisation may have been applied that cannot be reversed by division with a constant. If *ŝ*≠ 1, it is assumed that a linear normalisation was applied. In this case, the reciprocal of the sample-specific scaling factor is used to calculate the reversed normalised signal *ŷ* = *ŷ*_1_ ···*ŷ*_*N*_ (Eq. 23).

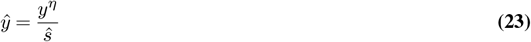

To correct for floating point errors, values in *ŷ* are rounded to the nearest integer and saved to a bigWig file. Assuming *ŝ* was estimated within a small margin of error, the original signal prior to linear normalisation can be accurately reconstructed.

